# Familiarity increases aggressiveness among clonal fish

**DOI:** 10.1101/386714

**Authors:** Carolina Doran, David Bierbach, Kate L. Laskowski

**Affiliations:** Department of Biology and Ecology of Fishes, Leibniz-Institute of Freshwater Ecology and Inland Fisheries, Berlin, Germany.

**Keywords:** Familiarity, Genetic relatedness, Amazon molly, clonal, group behaviour

## Abstract

Understanding how animal groups form and function is a major goal in behavioural ecology. Both genetic relatedness and familiarity among group mates have been shown to be key mediators of group composition. However, disentangling the two in most species is challenging as the most familiar individuals are often the most related, and vice versa. In order to gain a complete understanding of how individual interactions shape group behaviour it is crucial to understand the role each of this social relationship factors plays individually. To this end, we manipulated the level of familiarity among groups of the naturally clonal, and genetically identical fish, the Amazon molly (*Poecilia formosa*) and monitored group behaviour in an open-field and when given the opportunity to forage. Contrary to our predictions, fish that were the most familiar with each other showed the highest levels of aggression. Additionally, fish that were less familiar with each other exhibited the highest group cohesion and took the longest to begin feeding, compared to the more familiar fish. These results suggest that familiarity may socially buffer individuals from the perception of risk in novel environments, such as is common in most behavioural tests designed to test group behaviour. Increases in aggression that are associated with increasing familiarity as shown here might be a mechanism by which fish maintain a fission-fusion society with important consequences for the patterns of associations in group living animals.

## Introduction

Animals living in groups must balance the potential benefits of group living, such as greater foraging efficiency, against the costs, such as increased aggression and competition (Hamilton, 1971; Krause & Ruxton, 2002). For example, fish in shoals benefit from being able to better sense their local environment and have a higher probability of encountering food, proportional to the size of the group (Berdahl, Torney, Ioannou, Faria, & Couzin, 2013; Magurran & Pitcher, 1987; Peuhkuri, 1997). However group living also incurs costs, such as competition over resources. Thus there is a trade-off between costs and benefits of group living, particularly when it comes to food exploitation (Conradt & Roper, 2005; Ranta & Lindström, 1990; Reader, 2014). Individuals often mediate this trade-off by preferentially associating with related and/or familiar individuals (Lee-Jenkins & Godin, 2013; Thünken, Hesse, Bakker, & Baldauf, 2016; Ward & Hart, 2003). However, this can be complicated by the fact that animal groups often experience high group turnover meaning individuals have to constantly adjust their behaviour to that of new members (Couzin, 2006).

Groups are defined by individual animals aggregating and the patterns of interactions among these individuals can vary in response to several factors. Fish, for instance are known to alter their nearest neighbour distance in response to the presence of a predator or food (Hoare, Krause, Peuhkuri, & Godin, 2000; Sogard, 1997). Furthermore, fish often switch between shoals for any number of reasons, such as phenotypic assortment, differences in swimming speed or due to the existence of conflicting information (Croft et al., 2005; Killen, Marras, Nadler, & Domenici, 2017; Krause, Reeves, & Hoare, 1998; Merkle, Sigaud, & Fortin, 2015). In these fission-fusion societies, the patterns of social encounters among individuals are very dynamic, with important implications for the transmission of diseases and flow of information (Aureli et al., 2008; Couzin, 2006; Krause & Ruxton, 2002). These repeated associations will in turn affect the degree of familiarity among group members varying both among and within groups (Griffiths, 2003; Griffiths & Magurran, 1999; Lee-Jenkins & Godin, 2013; Utne-Palm & Hart, 2000; Ward & Hart, 2003). Understanding how and why individuals associate with each other can give insight into how groups may evolve and function. A major difficulty however to studying the effects of familiarity on group dynamics is that familiar individuals are often also more related, making disentangling these two critical factors difficult.

According to kin selection, genetic relatedness amongst individuals should facilitate cooperation and reduce competition and thus contribute to the evolution of group living (Hamilton, 1964). Genetic relatedness has been shown to increase cooperation and individual fitness (Gerlach, Hodgins-Davis, MacDonald, & Hannah, 2007; Hesse, Anaya-Rojas, Frommen, & Thünken, 2015; Thünken et al., 2016). Familiarity seems to add to the effects of kinship when it comes to group behaviour (Griffiths & Magurran, 1999; Lee-Jenkins & Godin, 2013). Several studies show how fish shoaling with familiar conspecifics tend to profit more from social learning, experience less aggression, find food faster and eat more than groups composed of unfamiliar individuals (Berdahl et al., 2013; Swaney, Kendal, Capon, Brown, & Laland, 2001; Utne-Palm & Hart, 2000). Associating with familiar individuals also appears to increase the anti-predator and foraging benefits of schooling behaviour (Chivers, Brown, & Smith, 1995; Metcalfe & Thomson, 1995; Pitcher, Magurran, & Edwards, 1985; Utne-Palm & Hart, 2000). However, the ecological context in which a group is, will influence whether both familiarity and/or kinship are beneficial or not (Frommen et al., 2012; Kelley, Graves, & Magurran, 1999; West, Pen, & Griffin, 2002).

A major step towards understanding the formation and function of groups relies on the ability to successfully disentangle familiarity and kinship which can be challenging in many species (Lee-Jenkins & Godin, 2013). Here we take advantage of a unique species to do just this. We used the naturally clonal fish, the Amazon molly (*Poecilia formosa*), a small live-bearing freshwater fish that produces broods of genetically identical offspring allowing us to isolate and test the role of familiarity on group dynamics. There is reason to believe that familiarity plays a major role in the group dynamics of the Amazon molly. These social fish form large groups in the wild where they forage and compete for resources together (Ingo Schlupp, Parzefall, & Schartl, 2002). These fish also strongly prefer to associate with familiar individuals (Makowicz, Tiedemann, Steele, & Schlupp, 2016; Ingo Schlupp, 2009); however it is currently unknown how the level of familiarity within a group influences group behaviour.

We predicted that fish that were more familiar with each other would show a decrease in aggression and resource defence (i.e. more egalitarian distribution of feeding). Furthermore we also predicted that this pattern would be accompanied by an increase in group cohesion.

## Methods

### The Amazon molly

This live-bearing, freshwater fish originated through a hybridization event between the Sailfin molly (*Poecilia latipinna*) and the Atlantic molly (*Poecilia. mexicana*) an estimated 100,000 years ago (Warren et al., 2018) and now reproduces through gynogenesis (Parzefall, 1989). This means that females require sperm from males of one of their parental species in order to induce embryogenesis, without incorporating its genetic content (Schartl, Wilde, Schlupp, & Parzefall, 1995).

### Fish husbandry

Fish were maintained in 100L stock tanks and fed ad libitum two times daily on standard flake food. Tanks were cleaned and 50% of the water volume was exchanged weekly. The population of *P. formosa* used for this study were obtained from Manfred Schartl, University of Würzburg. Since Amazon mollies reproduce gynogenetically and females require the sperm from one of their parental species in order to induce embryogenesis each stock tank also contained several male *P. mexicana*. We used a strain of *P. formosa* that has been kept in captivity since 2002 and regular molecular checks confirm that individuals are clones (M. Schartl. *personal communication*)

### Familiarity manipulation & group behaviour assays

We generated three treatment groups with mollies that had lived together for differing amounts of time. After this familiarity manipulation we measured their behavior in two different contexts: both before and after the addition of a defendable food resource.

We collected 96 individuals from 3 different stock tanks, assembling 24 groups of 4 individuals (8 groups per each treatment) in smaller housing tanks (40×20×20 cm). These groups were divided into 3 treatments – Low, Mid and High familiarity that differed in the amount of time individuals within each group spent together before having their behaviour observed – 0, 1 or 3 weeks respectively (Figure 1). To assemble an experimental group from the low and mid familiarity treatment groups, one individual from each of the 4 housing tanks were haphazardly removed and placed together. This was done either the day before behavioural observations for the low familiarity fish, or one week before for the mid familiarity fish. High familiarity fish were kept together for the entire three weeks. All fish were handled at the same time points as the other treatments were being assembled (e.g. low familiarity fish were handled at week 2 to mimic the assembling process being done in the mid familiarity fish). This involved all fish in the housing tank being netted, and held out of the tank for several seconds and then returned to their tank. This process ensured that all groups received the same amount and schedule of handling regardless of treatment.

**Figure 1.**
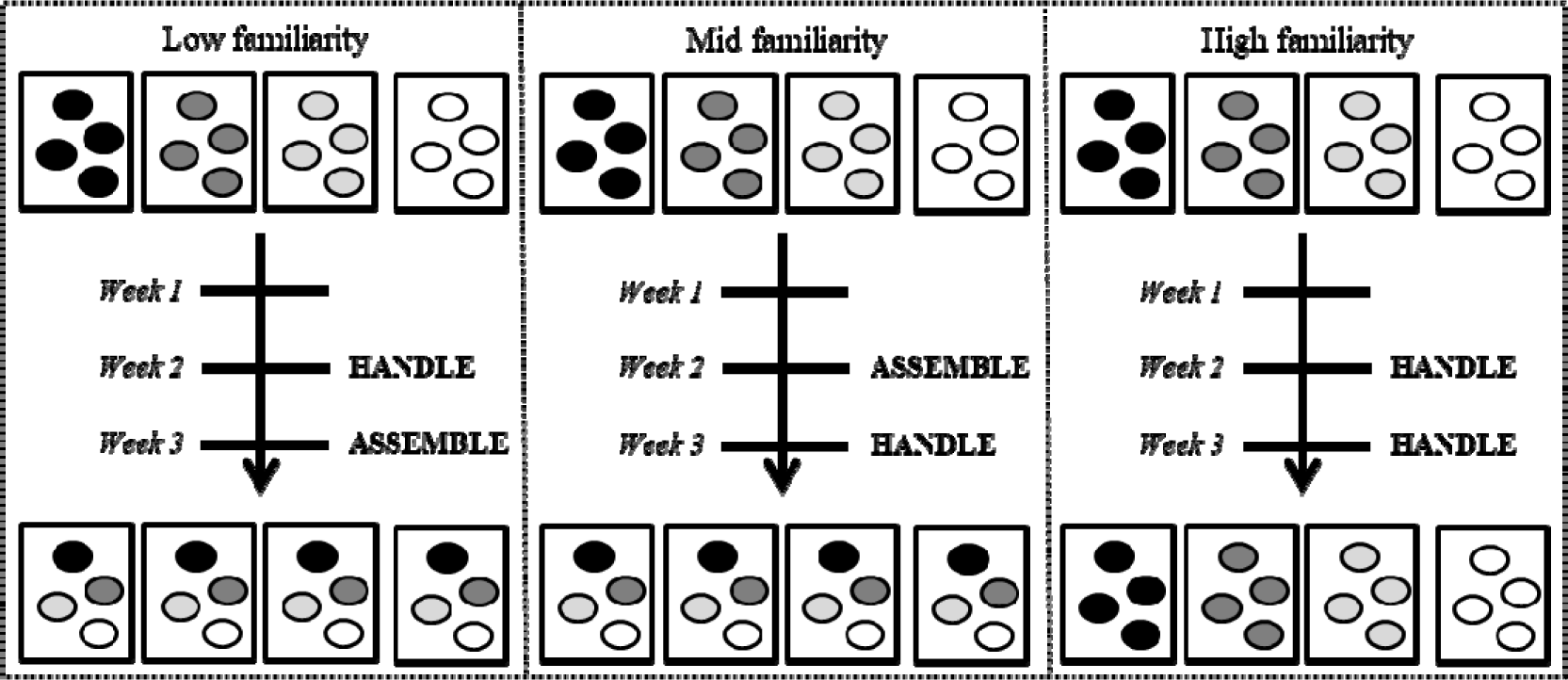
Detailed schedule of experimental procedure. Low familiarity groups were handled on week 2 and assembled the day before testing; Mid familiarity groups were assembled on week 2 and handled once more the day before testing. High familiarity groups were handled twice before being tested.

We started with 24 groups of 4 individuals (8 per treatment) but due to a video malfunction we lost the data corresponding to a group (Mid familiarity treatment). Finally due to natural mortality there were 3 groups with 3 individuals (1 in the high familiarity and 2 in the low familiarity treatment). Our final sample size was 20 groups of 4 individuals and 3 groups of 3 individuals.

After this familiarization phase, groups were transferred into an open field observation arena (60×30×30 cm) with 5cm deep water and allowed to acclimate overnight. The following morning, we observed the baseline behaviour of the fish in the observation arena for 5 minutes. After this time, we added a defensible food resource (sinking food tablet) and then observed the group for an additional 5 minutes after at least one individual had started feeding at the tablet. We counted the total number of aggressive interactions (bites, chases and tail beats) performed among group members (Bierbach et al., 2012; Foran & Ryan, 1994; Heubel & Plath, 2008), and the median inter-individual distance among group members (extracted from tracking paths from Ethovision Xt12, Noldus Information Technology, Inc) both before and after the addition of the food. After the food was added, we measured a number of foraging behaviours: latency for the first individual to begin feeding, how long each individual spent eating and the distribution of how many individuals were feeding simultaneously (i.e. one, two, three or four individuals at the food tablet at the same time). We additionally identified the dominant individual in each group as the individual who performed the majority of aggressive encounters while receiving the fewest (Bierbach et al., 2014; Laskowski, Wolf, & Bierbach, 2016). After having identified the dominant individual we also scored the amount of time it spent eating. Finally the collective duration of feeding was obtained by summing the duration of feeding by every individual in the group. At the end of every trial a snapshot form every video was used to measure the standard length of every individual.

### Statistics

We tested how familiarity and context (before and after food) influenced group behaviour using general linear mixed models. We ran separate models for the total number of aggressive behaviours and inter-individual distance. Both models included the fixed effects of familiarity (low, mid, high) and context (before/after the food was added) and an interaction between the two. We also included median body size as a covariate and group as a random effect. To investigate how familiarity influenced feeding behaviour, we used general linear models with latency to begin feeding and collective feeding as our response variables. These models included familiarity as a fixed effect and median group body size as a covariate; group was not included as a random effect as these response variables only had one observation per group. To test the overall significance of the fixed effects we compared the log-likelihood ratio of a model that contained the effect of interest, to one that did not. In order to obtain the amount of variation explained by each model we additionally estimated both the marginal and conditional R-squared value according to (Nakagawa & Schielzeth, 2013). These tell us how much variation was explained by the fixed factors or by both fixed and random factors together, respectively. Finally in order to test for differences in the number of animals feeding simultaneously we ran a chi-square test with the number of seconds there were one, two three or four individuals eating (for this test only groups of 4 individuals were counted). All statistics were run in R v3.4.4, using packages nlme and lme4 for linear and mixed models and ggplots and ggplot2 for plotting (Crawley, 2012). For all the models the residuals did not vary significantly from a normal distribution and there was no evidence of variance heterogeneity (see figures in SM). The R script of the entire statistical analysis can be found in the supplementary material.

### Ethical Note

This research was conducted in accordance with the ASAB/ABS guidelines for the use of Animals in Research. The reported experiments comply with current German law approved by LaGeSo Berlin (GO124/14 to D.B.).

## Results

### Aggression

Contrary to our predictions, we found that aggressive interactions significantly increased with familiarity in a clonal fish (table 1, significant effect of familiarity). We also found that aggression was higher before the food was added compared to after (table 1, significant effect of context). Interestingly, in the mid-familiarity treatment, the decrease in aggression after the food was added appears to be the strongest compared to the other two familiarity treatments (Figure 2), though this effect was not strictly significant (table 1, non-significant interaction.)

**Figure 2.**
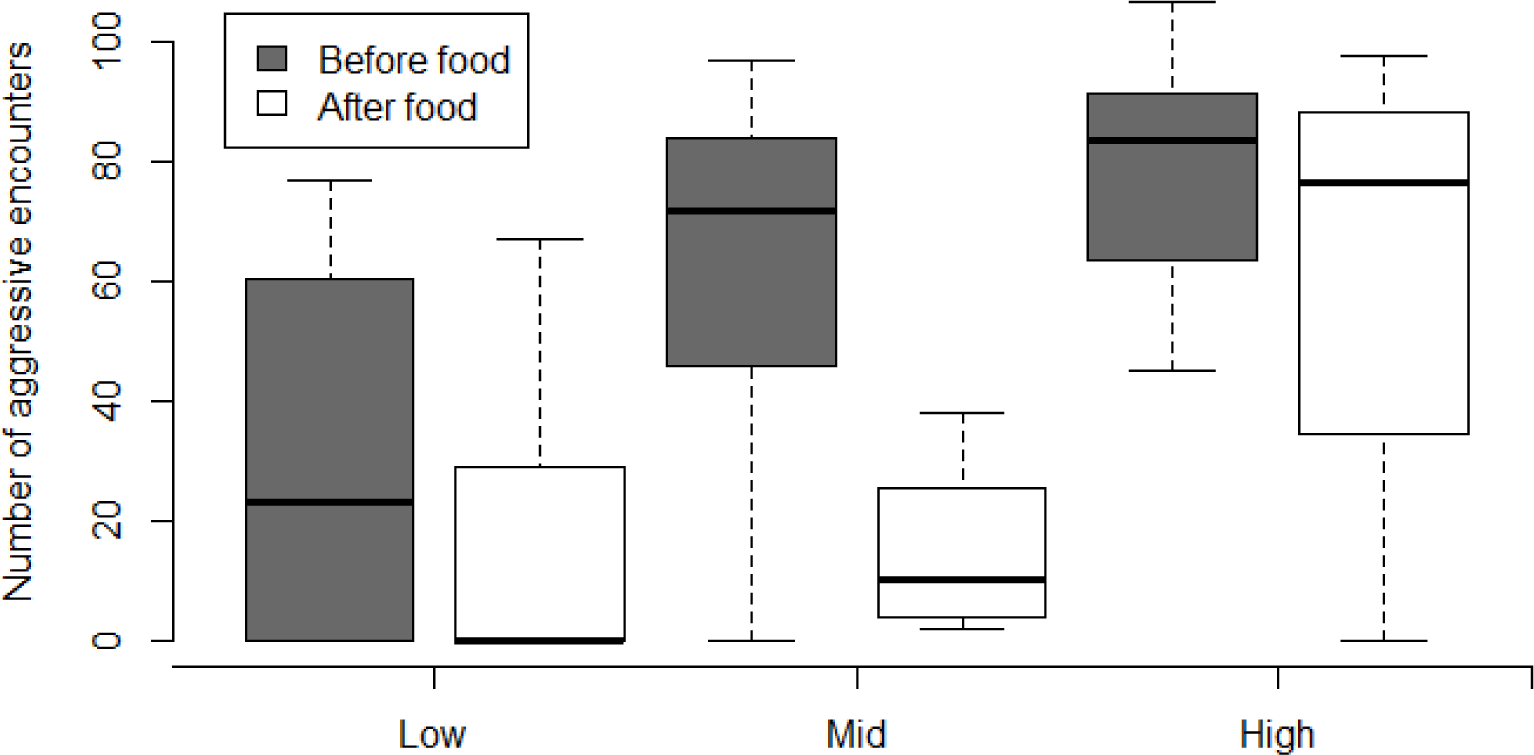
Boxplot of the total number of aggressive encounters among all members of a group within each treatment before (dark grey) and after (white) food addition. Dark lines represent medians, the box the inter-quartile range (IQR) and finally the whiskers are 2.5 times the IQR.

**Table 1.**
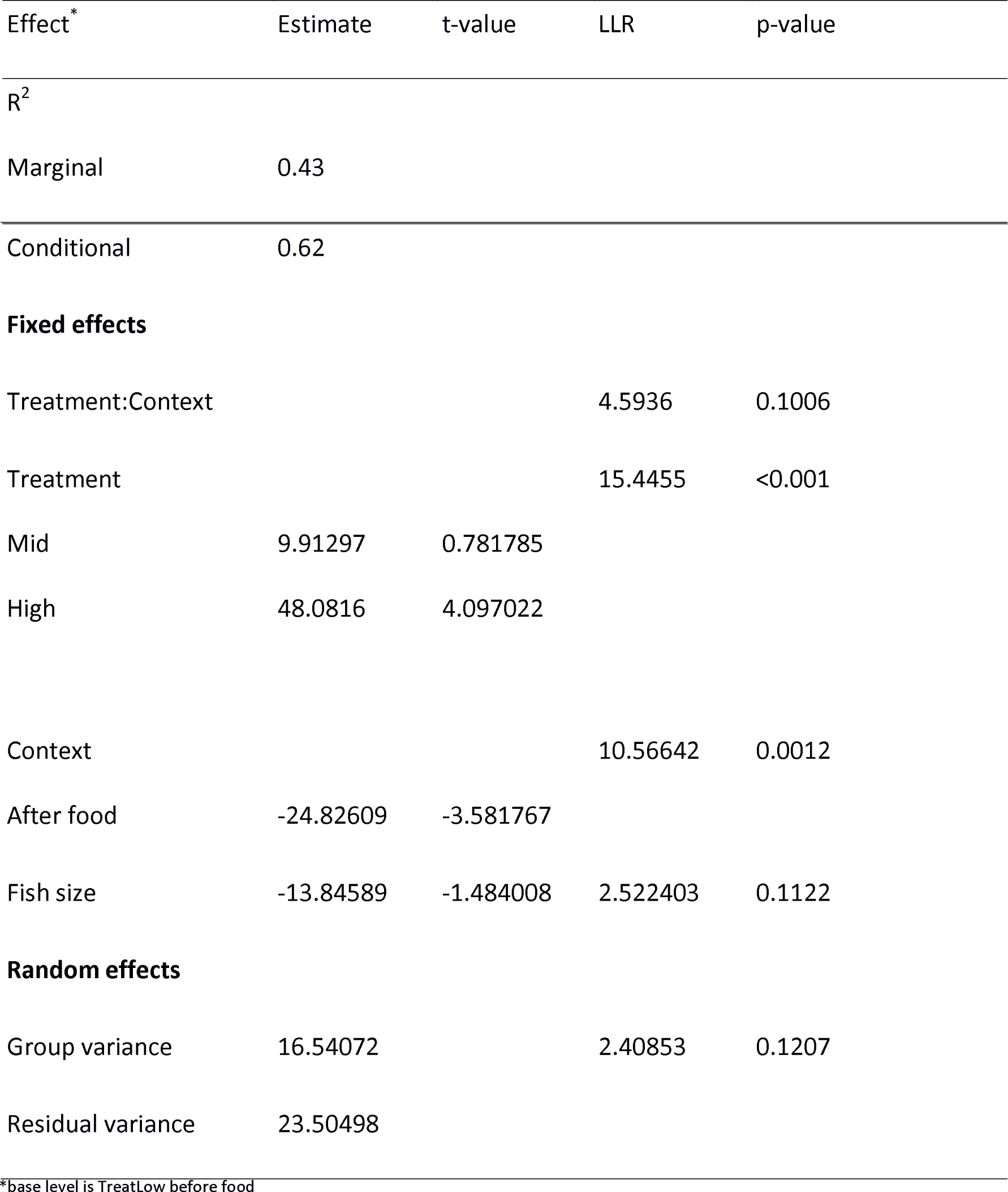
Results of the model testing the effects of treatment and context on total number of aggressive encounters within a group.

| Effect* | Estimate | t-value | LLR | p-value |
| --- | --- | --- | --- | --- |
| R<sup>2</sup> |  |  |  |  |
| Marginal | 0.43 |  |  |  |
| Conditional | 0.62 |  |  |  |
| **Fixed effects** |  |  |  |  |
| Treatment:Context |  |  | 4.5936 | 0.1006 |
| Treatment |  |  | 15.4455 | <0.001 |
| Mid | 9.91297 | 0.781785 |  |  |
| High | 48.0816 | 4.097022 |  |  |
| Context |  |  | 10.56642 | 0.0012 |
| After food | -24.82609 | -3.581767 |  |  |
| Fish size | -13.84589 | -1.484008 | 2.522403 | 0.1122 |
| **Random effects** |  |  |  |  |
| Group variance | 16.54072 |  | 2.40853 | 0.1207 |
| Residual variance | 23.50498 |  |  |  |

\*base level is TreatLow before food

### Feeding behaviour

Again, contrary to our predictions, the collective duration of feeding was not affected by familiarity (p-value = 0.1954, supplementary information). However, the time taken for the first individual to start feeding after the addition of food significantly decreased with familiarity (Table 2, significant effect of treatment). Furthermore, the distribution of feeding, i.e. how many individuals were eating at any given time within each group was significantly different for each of the treatments (X^2^ = 38087, df = 8 and p-value < 0.001). The amount of time the dominant fish of each group spent eating was not affected by treatment (p-value = 0.6267, supplementary information). What changed was how food was distributed within each group and in low familiarity, more individuals were able to simultaneously feed at the food tablet (Figure 3). Finally, body size did not significantly influence any measure of feeding behaviour.

**Table 2.** Results of the model testing the effects of treatment on latency to begin feeding within each group.

| Effect* | Estimate | t-value | f-value | p-value |
| --- | --- | --- | --- | --- |
| R<sup>2</sup> | 0.42 |  |  |  |
| Treatment |  |  | 5.8906 | 0.01 |
| Mid | -0.6405 | -2.033 |  |  |
| High | -0.9861 | -3.381 |  |  |
| Fish size | 0.293 | 1.264 | 1.596 | 0.2216 |

\*Base level is Treatment Low

**Fig 3.**
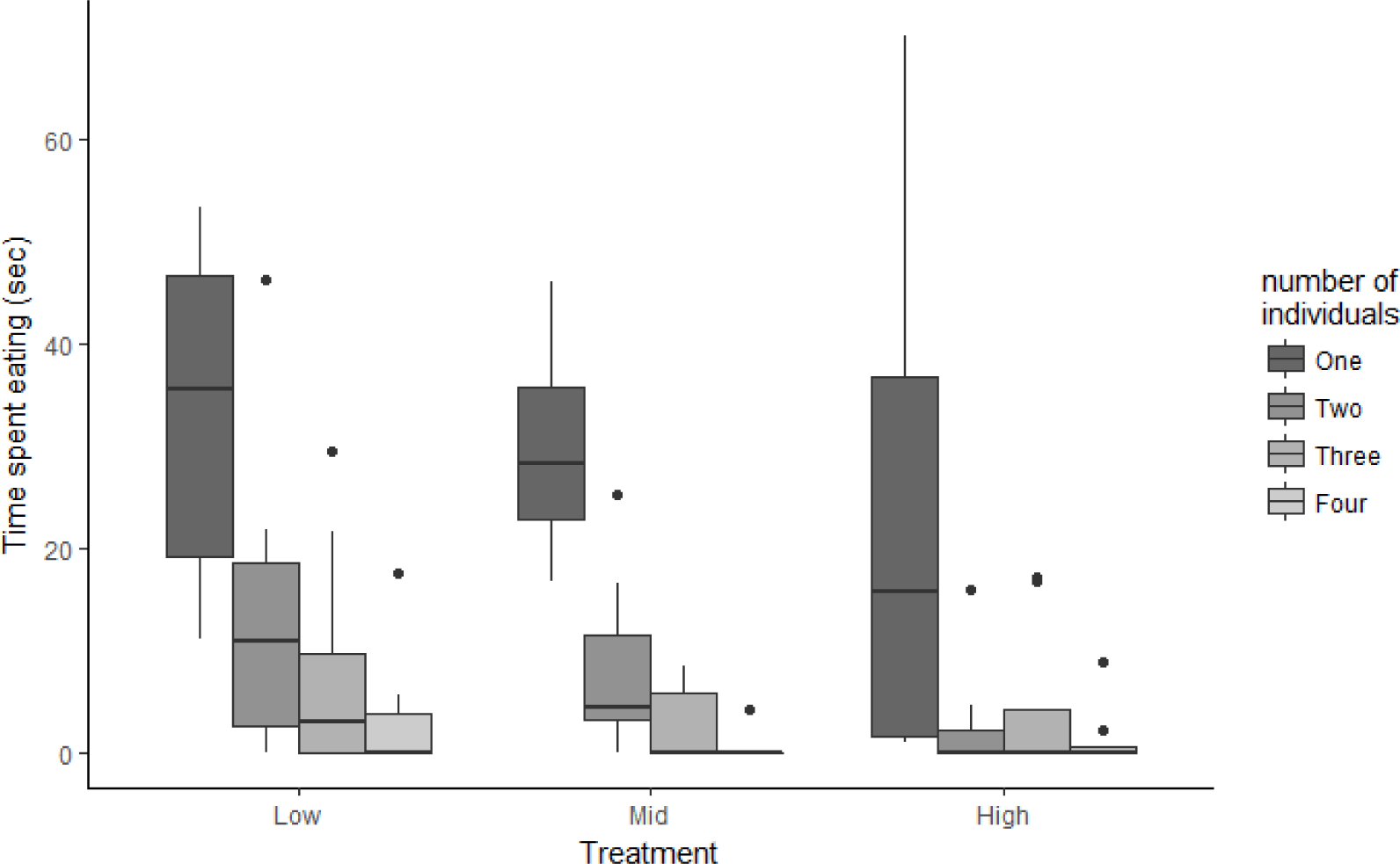
Amount of time one, two, three or four fish were observed feeding together in each treatment. Dark lines represent medians, the box the inter-quartile range (IQR) and finally the whiskers are 2.5 times the IQR.

### Group behaviour

Finally, also against our initial prediction, increasing familiarity within groups led to decreases in group cohesion as measured by inter-individual distance. Cohesion increased after food was added (table 3, significant effect of context) and this was independent of the familiarity treatment (non-significant interaction between treatment and context, Figure 4, Table 3). Body size had no effect.

**Fig 4.**
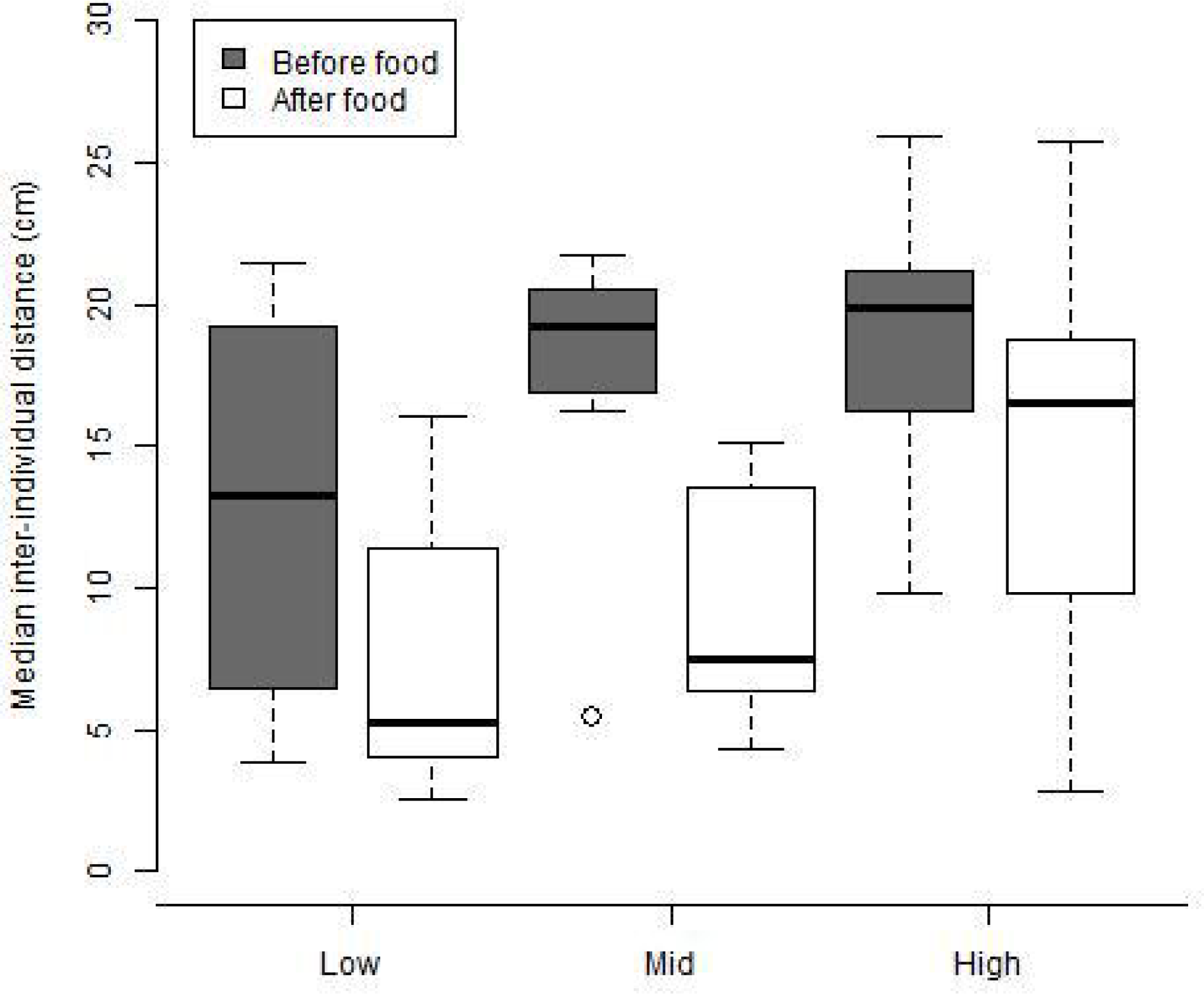
Median distance within individuals from a group Before Food (dark grey) and After Eating (white) for each treatment. Dark lines represent medians, the box the inter-quartile range (IQR) and finally the whiskers are 2.5 times the IQR.

**Table 3.** Results of the model testing the effects of treatment on median inter-individual distance within each group.

| Effect | Estimate | t-value | LLR | p-value |
| --- | --- | --- | --- | --- |
| $R^2$ | | | | |
| Marginal | 0.302 |  |  |  |
| Conditional | 0.593 |  |  |  |
| **Fixed effects** |  |  |  |  |
| Treatment:context |  |  | 1.5831 | 0.4531 |
| Treatment |  |  | 6.6621 | 0.0358 |
| Mid | 3.48 | 1.2785 |  |  |
| High | 6.57 | 2.5691 |  |  |
| Context |  |  | 13.46 | >0.001 |
| After food | -5.68 | 1.3721 |  |  |
| Fish size | 0.623987 | 0.3898 | 0.14637 | 0.702 |
| **Random effect** |  |  |  |  |
| Group variance | 3.928557 |  | 4.00942 | 0.0452 |
| Residual variance | 4.636645 |  |  |  |

## Discussion

A critical step towards understanding the formation of social associations in animal groups relies on the ability to disentangle the roles of familiarity and kinship. Kin selection tells us how genetic relatedness is at the core of the evolution of group living with several studies suggesting how familiarity adds to this effect (Griffiths & Magurran, 1999; Hamilton, 1964; Lee-Jenkins & Godin, 2013). But how does group behaviour differ based on familiarity alone? By exploiting a unique animal system, the clonal Amazon molly, we were able to isolate the effects of familiarity on the behaviour of fish groups. We show that, contrary to our predictions, familiarity significantly increased the number of aggressive encounters accompanied by a decrease in group cohesion.

This pattern contradicts previous studies showing that both for guppies and sticklebacks, being more familiar with individuals from their group not only decreases aggression as it also increases group cohesion and food sharing (Chivers et al., 1995; Höjesjö, Johnsson, Petersson, & Järvi, 1998; Johnsson, 1997). There are a number of potential explanations for this unexpected finding. Due to their unique type of reproduction this unisexual fish (*P. formosa*) lives in mixed-species shoals together with their parental species P. latipinna and P. mexicana, and also with different clonal lineages (I. Schlupp & Ryan, 1996). This renders their environment highly competitive in terms of mate choice and might explain why in this fish species aggression increases with familiarity. Previous research has already shown that *P. formosa* is more aggressive when compared to the parental species, and this aggression increases over time. (Makowicz & Schlupp, 2015) suggest that since in natural conditions, groups are very plastic with high group turnover, individuals might have low tolerance for consistent interactions with the same individuals. Furthermore, due to the fact that these fish require sperm from males of one of the parental species (Bierbach et al., 2011; Ingo Schlupp, Parzefall, & Schartl, 1991), Amazon mollies might be more aggressive to out-compete heterospecific females for access to males. This increase in aggression with familiarity may therefore provide one reason why so many fish species exist as dynamic fission-fusion societies.

Whereas familiarity had strong effects on aggression, there was no overall effect on familiarity in terms of the duration of time the group as a whole spent feeding. However, the distribution of food among the different individuals differed. The dominant individual was always the one who spent more time eating, and this was not affected by familiarity. When individuals were less familiar with each other, they each had more opportunity to access the food then when individuals were familiar, again contrary to what had been previously seen for other fish (Utne-Palm & Hart, 2000). This makes sense, as in these low familiarity groups aggression was lower and group members had more opportunity to reach the food. Thus individuals may be more motivated to frequently change groups (and decrease familiarity) if this increases their ability to access food resources.

Another potential explanation for why less familiar fish show unexpectedly low aggression is because of social buffering. Social buffering is the process by which social groups offer a safer environment in the presence of a perceived threat (Faustino, Tacão-Monteiro, & Oliveira, 2017). When familiarity is low being placed in a novel environment such as the testing arena used here might be perceived as risky. Thus individuals remain cohesive as a group with low levels of aggression. Indeed, after the addition of the food, which required a necessary disturbance, group cohesion in the low familiarity groups increased and individuals took the longest to begin feeding, further suggesting they interpret this disturbance as a potential threat. When individuals are more familiar with one another, as is the case for the mid familiarity treatment, the scenario changes slightly. Here, individuals exhibited lower group cohesion and higher aggression prior to the food being added, but once the group is disturbed by the addition of the food, aggression decreases and cohesion increases similar to what was seen in the low familiarity groups. Finally, three weeks of familiarization in the highly familiarity groups appears to be sufficient to eliminate any perceived risk by the introduction to the novel testing arena or the addition of the food as aggression remained high and group cohesion low in these groups. Taken together, our results suggest that familiarity mediates how the group responds to novelty, through social buffering. Groups that are more familiar can cope better both with being in a new environment and with the unexpected food delivery, and thus are able to maintain their high levels of aggression. The fact that aggression increases indicates that individuals are no longer seeking safety in numbers but instead asserting their dominance within their groups.

Both familiarity and genetic relatedness are known to be important factors for the development of a group’s social structure (Atton, Galef, Hoppitt, Webster, & Laland, 2014; Griffiths & Magurran, 1999). Working with genetically identical individuals provides the perfect opportunity disentangle the effects of both. In this study we were able to isolate the role of familiarity in modulating group behaviour. Understanding the factors that mediate group formation is at the forefront of studies of animal collective behaviour. That both familiarity and genetic relatedness are key is not new, but the contribution each plays separately is largely unknown. We were able to provide knowledge of how familiarity alone can contribute to the patterns of interactions within groups and their resulting foraging behaviour. Finally this study also sheds light into how aggression within groups may be critical for the maintenance of fission-fusion societies.

## Ackowledgements

We thank David Lewis, Marcus Ebert and Juliane Lukas for help with animal care. We are especially grateful to Hai Nguyen for his help with setting up the camera systems. Furthermore, we thank Manfred Schartl (University of Würzburg) for providing us with individuals of the Amazon molly. We received financial support from the Leibniz Competition (SAW-2013-IGB-2) and from the DFG (BI 1828/2-1 (to DB); LA 3778/1-1 (to KLL). Finally CD was supported by a AvH fellowship.

## Competing interests

The authors declare no competing interests.

